# Platelets as key cells in endometriosis patients: insights from small extracellular vesicles in peritoneal fluid and endometriotic lesions analysis

**DOI:** 10.1101/2024.08.21.607595

**Authors:** Barbara Bortot, Roberta Di Florio, Simona Merighi, Ben Peacock, Rebecca Lees, Francesco Valle, Marco Brucale, Alessandro Mangogna, Giovanni Di Lorenzo, Federico Romano, Gabriella Zito, Fabrizio Zanconati, Giuseppe Ricci, Valeria Cancila, Beatrice Belmonte, Stefania Biffi

**Affiliations:** Institute for Maternal and Child Health, IRCCS Burlo Garofolo, Via Dell’Istria 65/1, 34134 Trieste, Italy; Tumor Immunology Unit, Department of Health Promotion, Mother and Child Care, Internal Medicine and Medical Specialties “G. D’Alessandro”, University of Palermo, 90127, Palermo, Italy; NanoFCM Co., Ltd., Medicity Nottingham UK; Institute of Nanostructured Materials, Italian National Council of Research, Via Gobetti 101, 40129 Bologna, Italy; Department of Medical, Surgical and Health Science, University of Trieste, 34129 Trieste, Italy

**Keywords:** endometriosis, extracellular vesicles, platelet, neutrophil, follicular fluid, peritoneal fluid, GPIIb/IIIa, PF4

## Abstract

Endometriosis is a chronic inflammatory condition characterized by the presence of endometrium-like tissue outside the uterus, primarily affecting pelvic organs and tissues. In this study, we explored platelet activation in endometriosis. We utilized the STRING database to analyze the functional interactions among proteins previously identified in small extracellular vesicles (EVs) isolated from the peritoneal fluid of endometriosis patients and controls. The bioinformatic analysis indicated enriched signaling pathways related to platelet activation, hemostasis, and neutrophil degranulation. Double immunohistochemistry analysis for CD61 and MPO revealed a significant presence of neutrophils and platelets infiltrating endometriotic lesions, suggesting potential cell-cell interactions. Subsequently, we isolated small EVs from the peritoneal fluid of women diagnosed with endometriosis and from women without endometriosis who underwent surgery for non-inflammatory benign diseases. We performed single-particle phenotyping analysis based on platelet biomarkers GPIIb/IIIa and PF4 using nanoflow cytometry, as well as single-particle morphological and nanomechanical characterization through atomic force microscopy. The study demonstrated that patients with endometriosis had a notably higher proportion of particles testing positive for platelet biomarkers compared to the total number of EVs. This finding implies a potential role for platelets in the pathogenesis of endometriosis. Further research is necessary to delve into the mechanisms underlying this phenomenon and its implications for disease progression.

## 1. Introduction

Endometriosis is an estrogen-dependent chronic inflammatory process that affects 10– 15% of women of reproductive age ^1,2^. The condition is characterized by the presence of endometrium-like tissue outside the uterus, mainly in pelvic tissues, and can result in various types of debilitating pain and subfertility ^3^. Previous studies have shown that various alterations in inflammatory cells, complement components, cytokines, and chemokines within endometriotic lesions and peritoneal fluid contribute to the establishment of an inflammatory microenvironment ^4–7^. Significantly, this inflammatory niche interacts with endometriotic cells, including stromal and epithelial cells, playing a pivotal role in the development and persistence of endometriosis. For instance, the inflammatory microenvironment can enhance the survival and invasion of endometriotic cells ^8^ and contribute to the onset of pain and infertility associated with the condition ^9,10^. Moreover, the communication between inflammatory cells and endometriotic cells can perpetuate the inflammatory response, creating a harmful cycle that sustains the disease process ^11^. Moreover authors have demonstrated a link between the hemostasis system, specifically platelet hyperactivity, and the development and presentation of endometriosis symptoms ^12^. Platelets play a role in promoting angiogenesis, inflammation, and fibrosis within endometriosis lesions ^13–16^. Fibrosis is a biological process that involves activated platelets, macrophages, and myofibroblasts ^17^. Activated platelets in endometriosis lesions produce growth factors, cytokines, and chemokines such as TGF-β1, platelet-derived growth factor (PDGF), epidermal growth factor (EGF), and connective tissue growth factor (CTGF), which contribute to fibrogenesis ^17^. Research has shown a dynamic interaction between platelets and endometriotic lesions, leading to the development of endometriosis and causing a hypercoagulable state in affected women ^13,18^. Overall, a combined effect of the inflammatory response and hypercoagulation status may exist in advanced stages of endometriosis ^19,20^. Therefore, targeting the coagulation pathway through anticoagulant or antiplatelet therapy offers a potential treatment option for endometriosis and provides opportunities for identifying innovative biomarkers ^12,13^.

Extracellular vesicles (EVs) are membrane-bound particles that serve as highly efficient carriers of intercellular communication by transporting proteins, lipids, and nucleic acids ^21^. Studies have demonstrated that EVs play a crucial role in cellular communication in endometriosis by modulating immunosuppressive and proinflammatory pathways ^22,23^. Exploring EVs in endometriosis has the potential to offer valuable insights into the intricate mechanisms that drive the disease ^24,25^. Through cargo analysis, researchers have the potential to discover new biomarkers for early endometriosis detection or identify novel therapeutic targets ^26^. Moreover, deciphering how EVs contribute to the inflammatory microenvironment in endometriosis could pave the way for developing targeted treatments that disrupt this communication, potentially halting the disease”s progression.

Platelet-derived EVs are released during platelet activation ^27^ and can be measured using a variety of selectively expressed markers ^28^. In a previous study, we used single-particle phenotyping analysis based on platelet biomarkers, such as GPIIb/IIIa and PF4, to assess platelet activity in the ascites of women with ovarian cancer ^29^. Given the significant role of intraperitoneal fluid in the pathophysiology of endometriosis, as it serves as a dynamic interface between the immunological and reproductive systems ^4,5,30,31^, our goal was to evaluate platelet activity within this specific environment. Peritoneal fluid samples were collected from women diagnosed with endometriosis for this purpose. We employed the same single-particle phenotyping analysis that had successfully measured platelet activity in ovarian cancer patients.

## 2. Material and Methods

### 2.1. Proteome data enrichment and pathway analysis

The proteins of exosomes in peritoneal liquid from endometriosis patients and controls were retrieved from published literature in PubMed ^24^. The protein-protein interaction (PPI) network was visualized by the freely accessible, web-based Search Tool for Recurring Instances of Neighboring Genes (STRING) software ^32^. In addition to PPI network analysis, functional enrichment analysis using the Reactome database was performed.

### 2.2. Patient cohort and sample collection

Patients who underwent elective laparoscopy for suspected endometriosis due to pain symptoms and/or infertility were recruited during the preoperative assessment. Following approval by the Institutional Review Board (IRB-BURLO 01/2022, 09.02.2022), patients were asked to sign an informed consent form. The inclusion criteria for endometriosis patients were as follows: i) had a histological confirmation of superficial peritoneal endometriosis (SPE); ii) had a histological confirmation of deep infiltrating endometriosis (DIE); and iii) had a histological confirmation of ovarian endometrioma. The inclusion criteria for control patient selection were patients without endometriosis who underwent surgery for benign, not inflammatory, conditions. The exclusion criteria were as follows: i) interdicted patients who were unable to provide informed consent; ii) patients with other forms of endometriosis without endometriosis according to the inclusion criteria; iii) patients with peritoneal inflammatory diseases (i.e., pelvic inflammatory disease, diverticulosis, etc.); and iv) oncological patients with peritoneal involvement. The follicular fluid (FF) of the first and largest punctured follicle from the bilateral ovaries was collected during the oocyte retrieval procedure (transvaginal follicular aspiration). Each ovarian follicle was aspirated independently. As previously described ^29^, hemoglobin levels were measured in all the samples to determine the presence of blood contamination. Only uncontaminated samples were included.

Peritoneal fluid from 12 patients was used in the present study. The patients” characteristics and medications are reported in **Table 1**. We included 70 patients in the study, 11 of whom had free fluid in the pouch of Douglas. Six blood-free samples were used in this investigation. Additionally, follicular fluid from 5 infertile patients who underwent in vitro fertilization was analyzed.

**Table 1.**
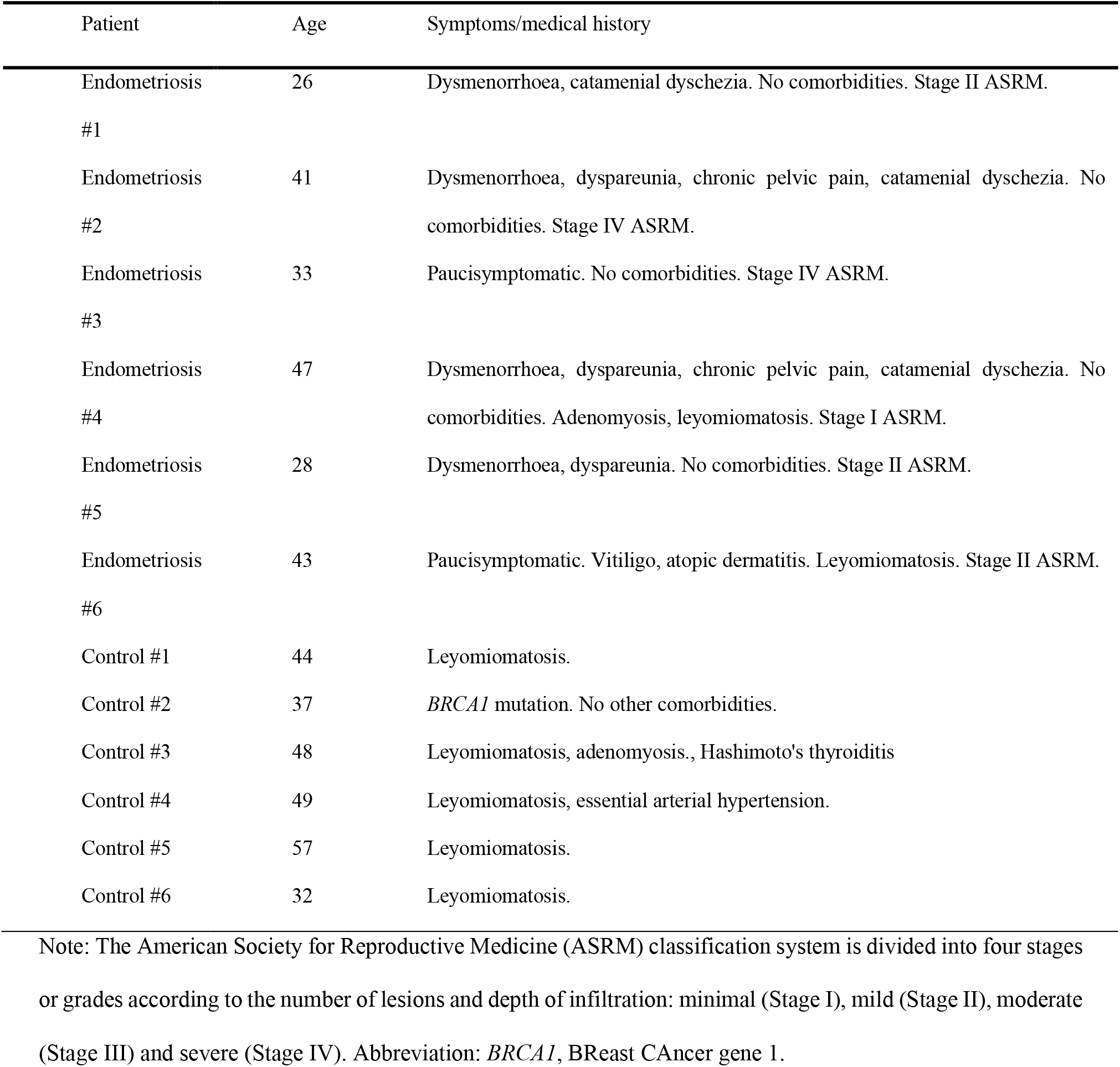
Patient characteristics.

| Patient | Age | Symptoms/medical history |
| --- | --- | --- |
| Endometriosis<br>#1 | 26 | Dysmenorrhoea, catamenial dyschezia. No comorbidities. Stage II ASRM. |
| Endometriosis<br>#2 | 41 | Dysmenorrhoea, dyspareunia, chronic pelvic pain, catamenial dyschezia. No comorbidities. Stage IV ASRM. |
| Endometriosis<br>#3 | 33 | Paucisymptomatic. No comorbidities. Stage IV ASRM. |
| Endometriosis<br>#4 | 47 | Dysmenorrhoea, dyspareunia, chronic pelvic pain, catamenial dyschezia. No comorbidities. Adenomyosis, leiomyomatosis. Stage I ASRM. |
| Endometriosis<br>#5 | 28 | Dysmenorrhoea, dyspareunia. No comorbidities. Stage II ASRM. |
| Endometriosis<br>#6 | 43 | Paucisymptomatic. Vitiligo, atopic dermatitis. Leiomyomatosis. Stage II ASRM. |
| Control #1 | 44 | Leiomyomatosis. |
| Control #2 | 37 | *BRCA1* mutation. No other comorbidities. |
| Control #3 | 48 | Leiomyomatosis, adenomyosis., Hashimoto's thyroiditis |
| Control #4 | 49 | Leiomyomatosis, essential arterial hypertension. |
| Control #5 | 57 | Leiomyomatosis. |
| Control #6 | 32 | Leiomyomatosis. |

Note: The American Society for Reproductive Medicine (ASRM) classification system is divided into four stages or grades according to the number of lesions and depth of infiltration: minimal (Stage I), mild (Stage II), moderate (Stage III) and severe (Stage IV). Abbreviation: *BRCA1*, BReast CAncer gene 1.

### 2.3. Small-EV isolation

The isolation of small EVs from peritoneal and follicular fluids was conducted according to a previously established method ^33^ using Total Exosome Isolation Reagent (Invitrogen, CN 4484453) and according to the manufacturer”s provided methodology. Briefly, we centrifuged the samples at 2000 × g to remove cells and debris. Subsequently, we recovered the small EVs from the resulting liquid through precipitation.

### 2.4. Nanoflow cytometry

The nanoflow cytometry (nFCM) procedure was carried out using a previously established method ^29^. Briefly, 10 µl of the EV sample was incubated with 2e^10^ particles/ml (PF4: 1:10, GPIIb/IIIa: 1:10, CellMask: 1:500) before incubation at room temperature for 1 h. After incubation, the mixture was diluted in PBS to 1 ml and ultracentrifuged for 1 hour at 100,000 × g using a Beckman Coulter benchtop optima TLX. For rapid nFCM analysis, pellets were resuspended to 1e^8^-1e^9^ particles/ml after supernatant removal. The nFCM Professional Suite v1.8 program processed the data and generated dot plots, histograms, and statistical data in a PDF. Gating inside the program enabled proportional analysis of subpopulations segregated by fluorescent intensities, with size distribution and concentration accessible for each.

### 2.5. Atomic Force Microscopy (AFM)

AFM-based quantitative morphology was performed as described elsewhere to obtain the size and stiffness distributions of the individual particles found in all the samples ^34^. Briefly, 5 µl aliquots from EV samples were deposited on poly-L-lysine-functionalized glass coverslips and left to adsorb for 30 min at 4°C. AFM micrographs were recorded in liquid at RT on a Multimode 8 microscope (Bruker, USA) equipped with Scanasyst Fluid+ probes (Bruker, USA) and operated in peak-force mode. Image analysis was performed with Gwyddion v2.61 and custom Python scripts ^35^. All samples were prepared following the same deposition protocol, thus allowing us to estimate the relative particle concentrations of the samples via their respective particle surface densities, as discussed elsewhere ^36^.

### 2.6. Immunohistochemical Analysis

Endometriosis tissue samples were fixed in a 10% v/v buffered formalin solution before being embedded in paraffin (formalin-fixed and paraffin-embedded, FFPE). Tissue sections, cut to a thickness of four micrometers, were then deparaffinized and rehydrated. The antigen unmasking technique was performed, using citrate buffer, pH 6, or Tris-EDTA buffer, pH 9, in thermostatic bath at 98°C for 30 min. Sections were then allowed to reach room temperature and washed in phosphate-buffered saline (PBS). After neutralization of endogenous peroxidase activity with 3% v/v H_2_O_2_ and blocking of non-specific bindings by PBS + 2% bovine serum albumin, samples were incubated with CD61 primary antibodies overnight at 4°C. Staining was revealed via anti-rabbit HRP-conjugated secondary antibody and 3,3′-diaminobenzidine (DAB, Dako, Agilent, Denmark) substrate chromogen. Slides were then counterstained with Mayer Hematoxylin (Diapath, Italy).

Multiplex immunohistochemical staining was performed on FFPE human tissue sections as previously described. Briefly, three-micrometer-thick tissue sections were deparaffined, rehydrated and unmasked using Novocastra Epitope Retrieval Solutions pH9 in a thermostatic bath at 98°C for 30 minutes. The sections were brought to room temperature and washed in PBS. After neutralization of endogenous peroxidase with 3% H_2_O_2_ and Fc blocking with 0.4% casein in PBS (Leica Biosystems), the sections were incubated with the following primary antibodies: mouse monoclonal CD61 (ready to use, clone 2F2 cod. PA0308, Leica Biosystems) and rabbit polyclonal MPO (myeloperoxidase) (1:25, cod. ab9535, Abcam). Double IHC staining was performed by applying SignalStain^®^ Boost IHC Detection mouse, horseradish peroxidase (HRP) conjugated produced in goat (cod. 8125S, Cell Signaling Technology) and SignalStain^®^ Boost IHC Detection rabbit, alkaline phosphatase (AP)-conjugated produced in horse (cod. 18653S, Cell Signaling Technology) and DAB (3,3′-Diaminobenzidine, Leica Biosystems) or Vulcan Fast Red (BioOptica) as substrate chromogens. All the sections were analyzed under Zeiss Axio Scope A1, and microphotographs were acquired using Axiocam 503 Color digital camera with the ZEN2 imaging software.

## 3. Results

Nazry *et al*. identified a set of unique proteins derived from exosomes isolated from the peritoneal fluids of endometriosis patients and controls ^24^. We employed bioinformatic data analysis to characterize signaling dynamics, and the PPI network analysis findings confirmed extensive and intricate interactions within the unique protein lists. **Figure 1** depicts the predicted PPI network during the proliferative phase of control and endometriosis stages I and II. Nodes represent proteins, edges indicate functional and physical protein associations, line thickness reflects the strength of data support, and only interactions with a high confidence score of ≥ 0.7 are visualized. The pathways of the innate immune system, neutrophil degranulation, platelet degranulation, hemostasis, and formation of the cornified envelope were identified as the top pathways that comprised an interconnected set of proteins in all of the samples analyzed (**Supplementary 1** shows the analysis of controls and endometriosis stage I/II in the secretive phase).

**Figure 1.**
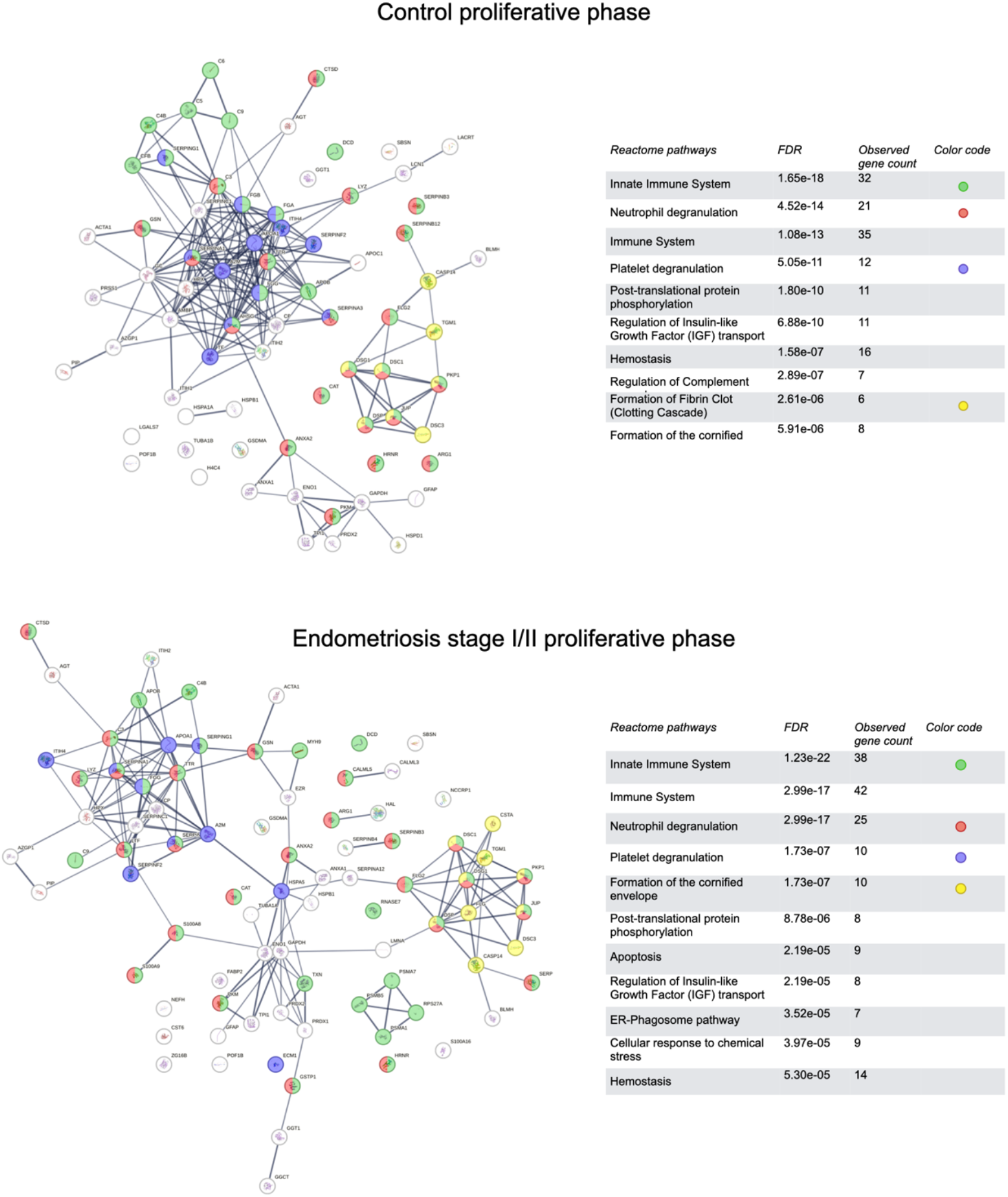
STRING protein-protein interaction (PPI) network analysis of proteins found in exosomes isolated from the peritoneal fluid during the proliferative phase in control patients (upper panel) and endometriosis patients (lower panel). In this network, nodes represent proteins, edges indicate functional and physical protein associations, and the thickness of the lines reflects the strength of data support. Only interactions with a high confidence score of ≥ 0.7 are displayed. The functional pathways were obtained from the Reactome database, and the top 10 pathways are listed in ascending order based on their *p*-value. FDR: False Discovery Rate.

Subsequently, we isolated small EVs from the peritoneal fluid of women diagnosed with endometriosis and from women without endometriosis who underwent surgery for non-inflammatory benign diseases. **Figure 2** illustrates the workflow of the analyses performed on the samples.

**Figure 2.**
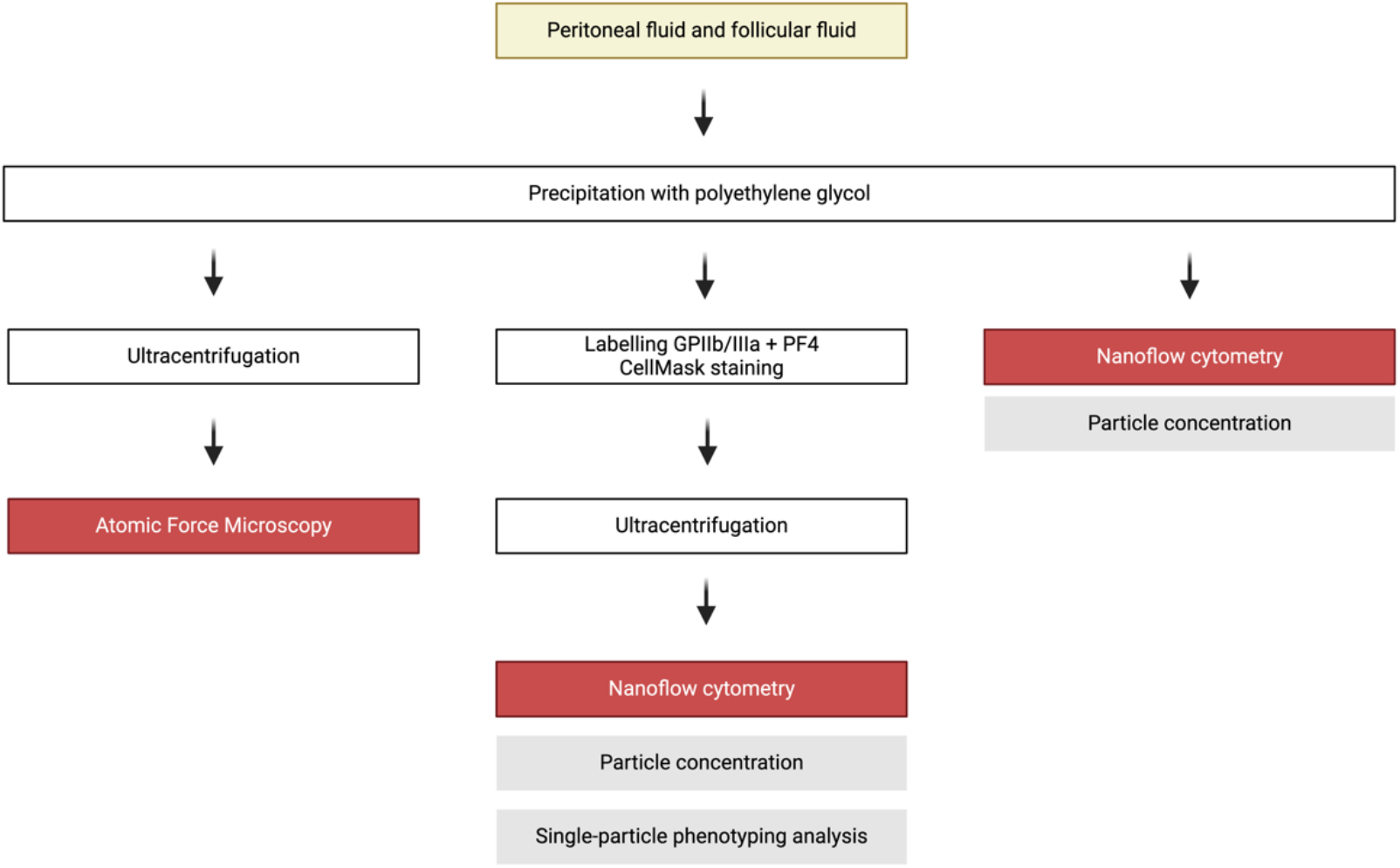
Overview of the analytical workflow.

The precipitation isolated samples contained approximately 10^9^ to 10^11^ particles/ml, measured via FCM, which decreased to 10^8^-10^9^ after UC. Both nFCM and AFM agreed to assign the highest particle concentrations to follicular fluid samples after UC. However, these follicular fluid-isolated particles showed low proportions of Cell Mask Deep Red (CMDR) labeling, typically below 20%. Little difference was observed between the control and endometriosis particle concentrations post-UC and the CMDR labeling, which, on average, accounted for 40% of the particles in these samples. The characteristics of the particle concentration, size, and percentage of membrane-positive particles in the samples measured via nFCM are shown in **Figure 3**. The combined morphological and nanomechanical characteristics of the individual particles in the selected samples were measured via AFM, confirming abundant EVs and residual content of non-vesicular particles with diameters less than 30 nm (**Figure 4**). Particles displaying nanomechanical behavior previously associated with intact vesicles ^34,36^ were found to have an average contact angle of ∼90°, which corresponds to a mechanical stiffness largely in line with values previously measured with the same technique on, *e*.*g*., samples isolated from ascites ^29^.

**Figure 3.**
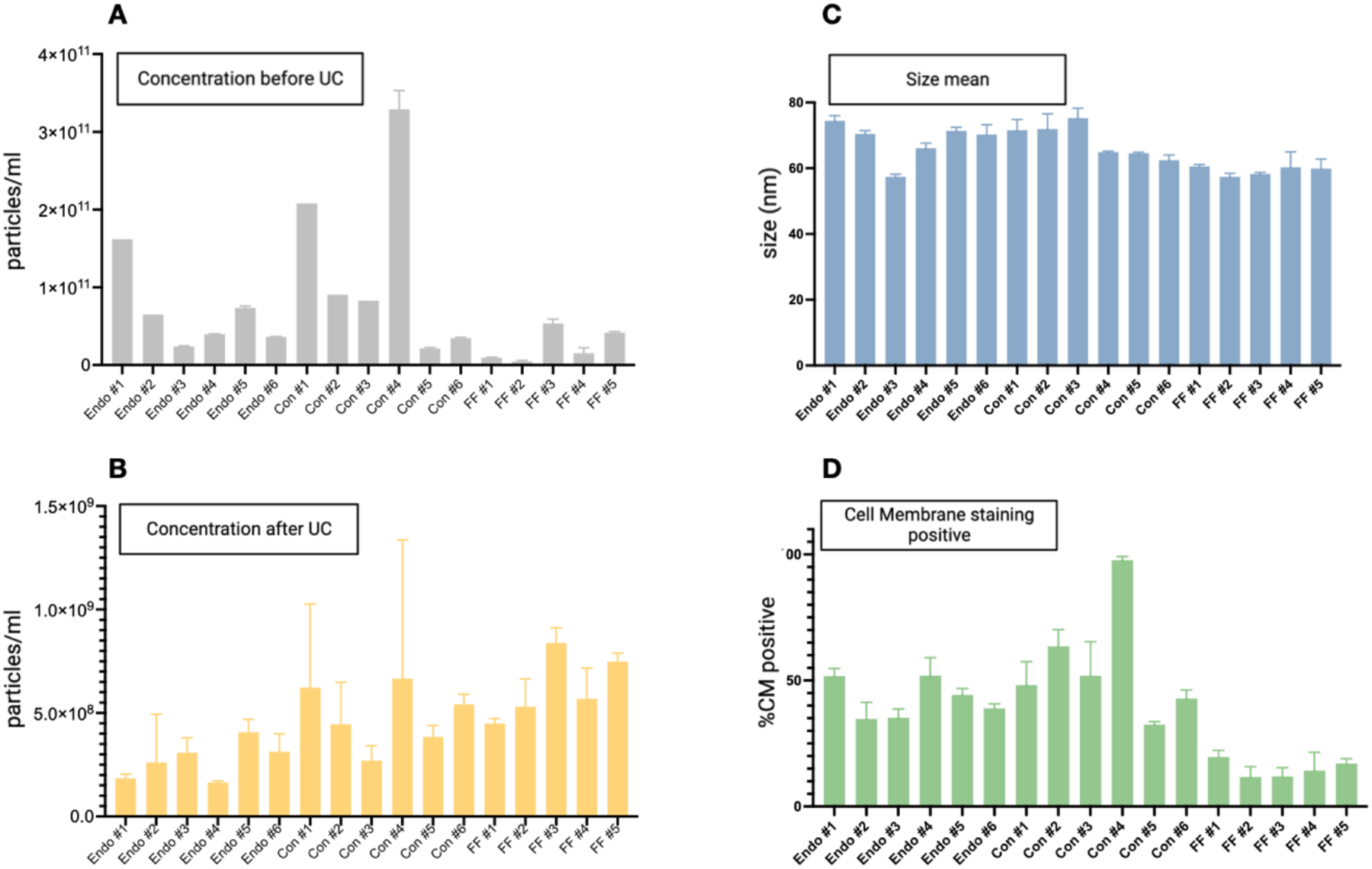
Nanoflow cytometry analysis (NanoFCM) of small EVs from peritoneal and follicular fluids in a cohort of patients. The data are reported as the means ± SDs of one experiment performed in triplicate. (**A**) The concentration of all particles > 45 nm in diameter in precipitation-isolated samples. (**B**) The concentration of all particles > 45 nm following ultracentrifugation (UC). (**C**) Size following UC. (**D**) The percentage of all particles > 45 nm, which also showed CellMask™ labeling (thus determined as EVs). Abbreviations: Endo, endometriosis sample; Con, control sample; FF, follicular fluid sample.

**Figure 4.**
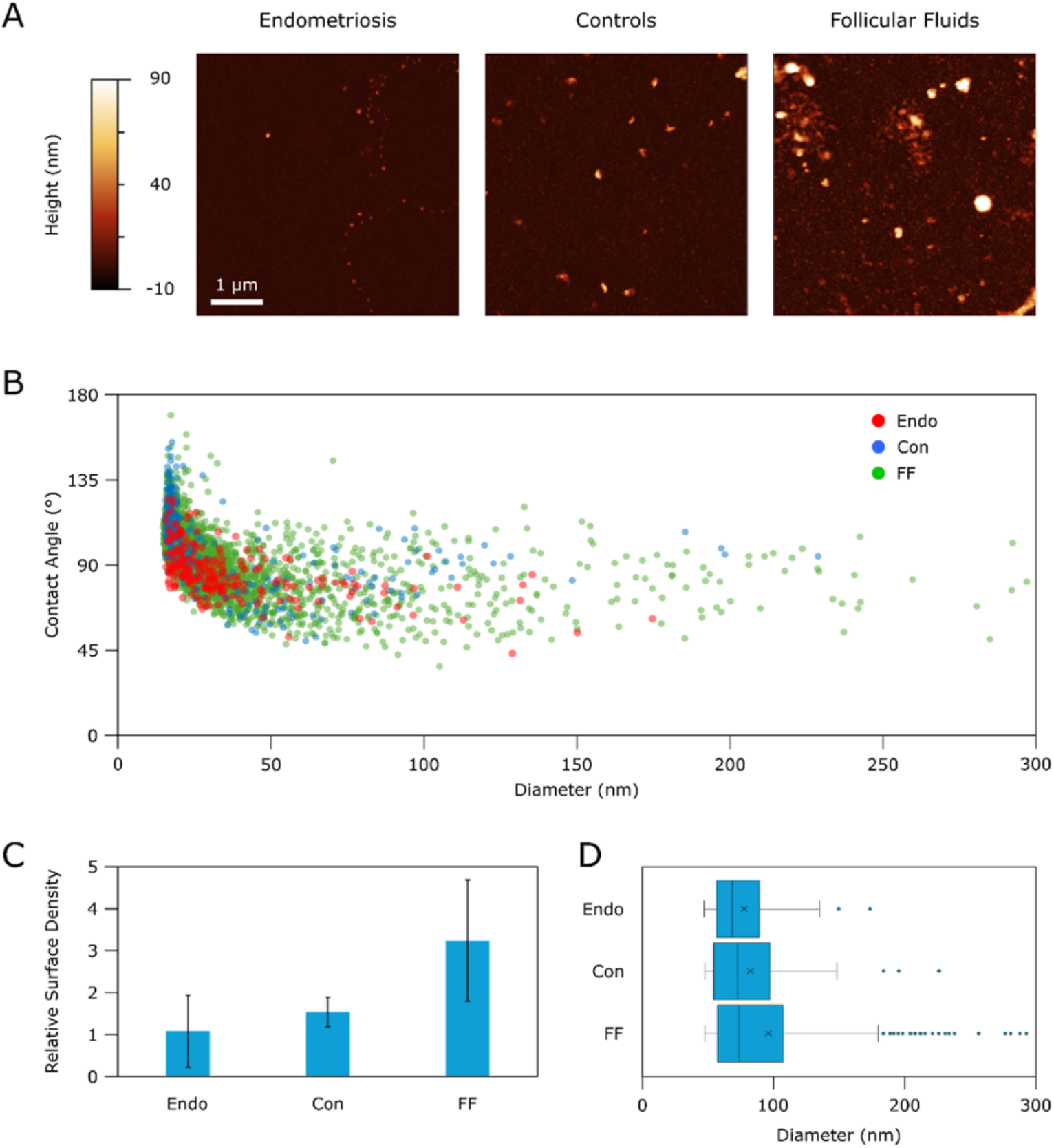
Atomic force microscopy (AFM)-based morphometry and nanomechanical screening. (**A**) Representative AFM micrographs of the three sample types. (**B**) Contact angle (CA) *vs* diameter plot of individual particles measured from at least 10 AFM micrographs deposited from duplicate preparations (Endo: N=227; Con: N=296; FF: N=3632). The contact angle can be regarded as a measure of the mechanical stiffness of EVs; all particles above ∼45 nm display the typical conserved average CA of intact vesicles, whereas smaller particles do not. (**C**) Normalized relative surface density of particles with diameters greater than 45 nm, showing that the FF samples are consistently more concentrated than the Endo and Con samples. (**D**) Box plot of the measured diameters of individual vesicles greater than 45 nm shows how FF samples contained the most significant proportion of larger EVs, while Endo samples contained the smallest. Abbreviations: Endo, endometriosis sample; Con, control sample; FF, follicular fluid sample.

Single-particle phenotyping analysis enabled the measurement of the platelet-associated markers GPIIa/IIIb and PF4 in all the samples, as well as EV membrane staining (**Figure 5**). The percentages of small-EVs that were positive for platelet markers in follicular fluids and control fluids were less than 2.9% and 3.7%, respectively, with no statistically significant differences between these two sets of samples. Endometriosis patients” samples had higher levels, ranging from 3.4% to 17%, with a statistically significant difference between these samples and follicular fluid and control samples.

**Figure 5.**
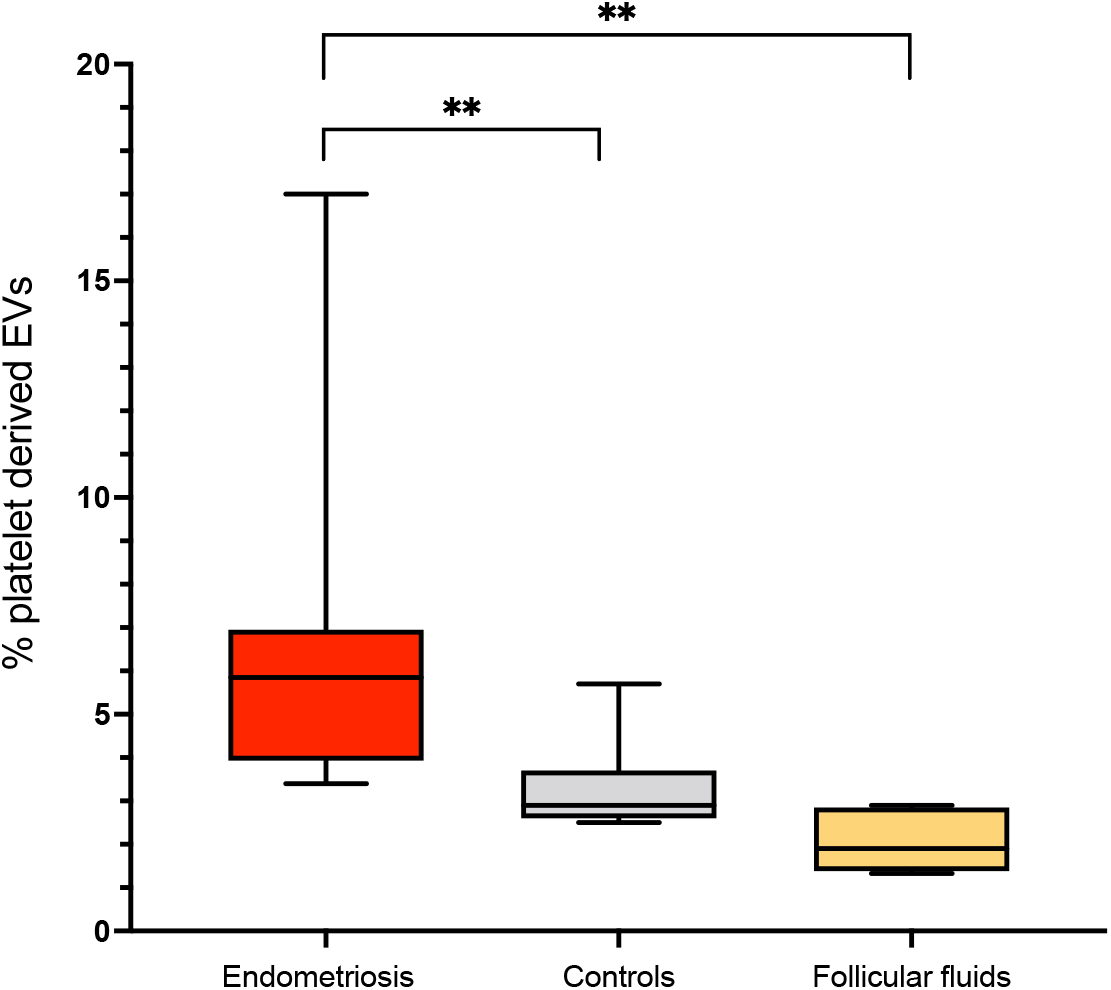
Percentage of particles positive for GPIIb/IIIa and PF4 relative to total EVs from peritoneal fluid in women with endometriosis (endometriosis), in women without endometriosis during surgery for a benign, non-inflammatory, condition (controls), and in follicular fluid samples collected from infertile patients undergoing in vitro fertilization (follicular fluids). The samples were fluorescently labeled with FITC-conjugated antibodies specific for GPIIb/IIIa and PF4. Additionally, the dye CellMask™ deep red plasma membrane stain (CM) was used to distinguish EVs from other small particles.

We performed immunohistochemistry (IHC) for CD61 on FFPE biopsies from patients with endometriosis to examine the presence and distribution of platelets. As shown in **Figure 6**, CD61-positive platelet clusters were observed surrounding the endometriotic lesions.

**Figure 6.**
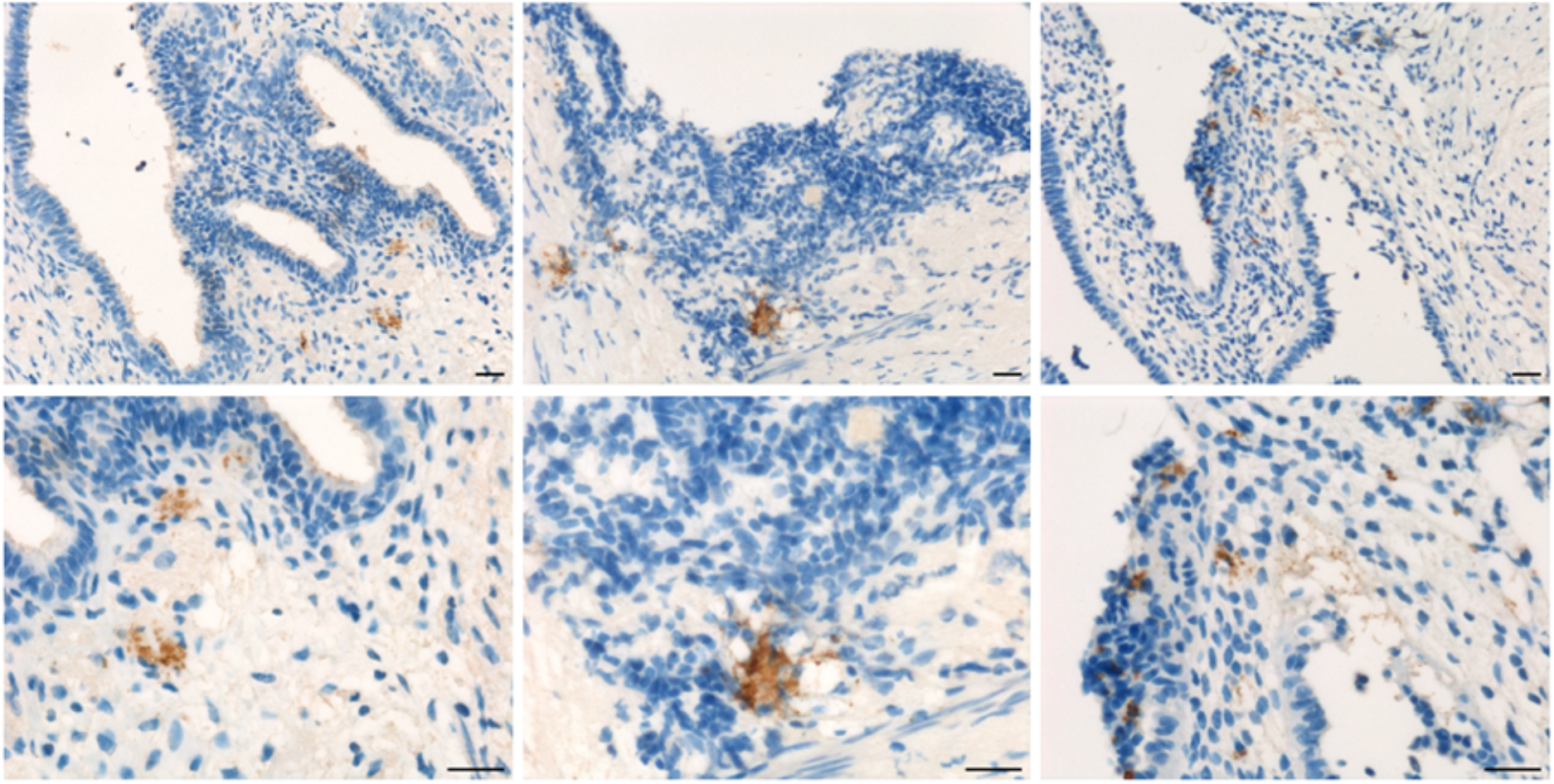
Microphotographs showing platelets in endometriotic lesions. Representative immunohistochemical staining of CD61. DAB (brown) chromogen was used to visualize the binding of the anti-human CD61 antibody. Magnifications were 200x (up), and 400x (down); scale bar, 50 µm.

We then investigated platelet activation within endometrioid lesions, focusing on their interaction with inflammatory cells using a multiplex immunolocalization assay for CD61 and MPO. Our analysis revealed both tissue and intravascular deposits of platelets within the lesions, with CD61-positive platelets closely associated with MPO-expressing granulocytes. This suggests a potential cell-cell interaction between neutrophils and platelets infiltrating the lesions (**Figure 7**).

**Figure 7.**
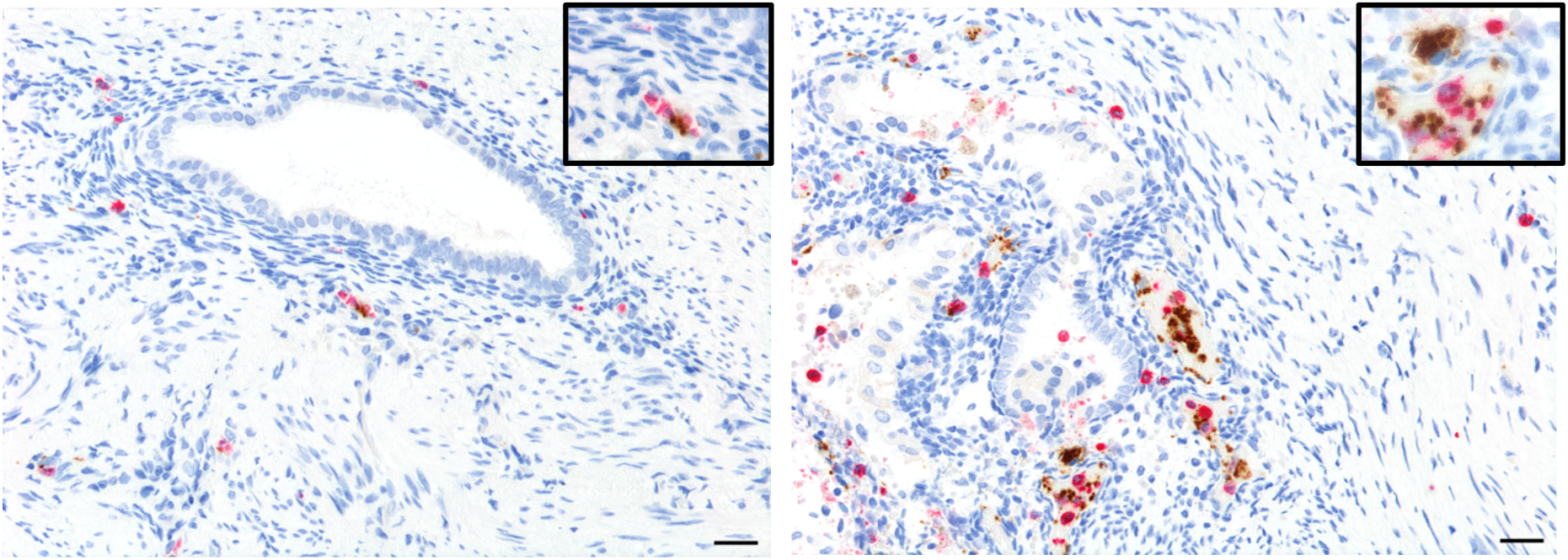
Representative microphotographs of double immunohistochemical staining for CD61 and MPO in endometriotic foci highlighting a close-contact between platelet aggregates (CD61+) and neutrophils (MPO+) in the cytogenic stroma of endometriotic lesions. As shown in inset at the bottom of the pictures, this interaction occurs at tissutal and intravascular sites. DAB (brown) chromogen was used to visualize the binding of the anti-human CD61 antibody, while AEC (red) chromogen was used to visualize the binding of the anti-human MPO antibody. The images were captured at magnifications of 200x (top) and 400x (bottom); scale bar, 50 µm.

## 4. Discussion

Endometriosis is considered a chronic inflammatory condition, and the primary approach to medical treatment involves the use of hormonal and anti-inflammatory medication ^2,37^. Scientific data shows a strong association between the coagulation system and the progression of endometriosis ^12,13,18,19^. Recent study data has shown the role of platelets in the progression of endometriosis, which supports the use of anticoagulants as a treatment for endometriosis. Additionally, it emphasizes the significance of evaluating platelet activation in individuals with endometriosis. Together with myofibroblasts, macrophages, and sensory nerve fibers, platelets are among the main cellular elements implicated in the pathophysiology of fibrosis ^17^.

Consistent with the literature ^13^, this research has found that platelet agglomerates in the cytogenic stroma of lesions suggest their migration from the microenvironment into the lesions. It was shown that platelets play essential roles in the development of endometriosis^13,14,38–40^ and it was found that endometriotic stromal cells release platelet-activating molecules^41^. We also demonstrated a possible interaction between platelets and neutrophils in endometriosis foci. Platelets release signals that attract neutrophils to sites of inflammation. For example, they secrete chemokines like CXCL4 (PF4) and CXCL7 that recruit neutrophils^42^. Platelets express P-selectin on their surface upon activation, which binds to PSGL-1 on neutrophils, facilitating their interaction ^43,44^. Platelets can induce the formation of neutrophil extracellular traps (NETs), which could play a role in the pathophysiological mechanisms of endometriosis ^45^. Both platelets and neutrophils release pro-inflammatory cytokines that amplify inflammatory responses.

To further understand this occurrence, we conducted a series of analyses to determine the presence of platelet activation in the peritoneal fluid of endometriosis patients. The bioinformatic analysis of proteins detected in exosomes of the intraperitoneal fluid has revealed specific biochemical pathways, such as those associated with the immune system, hemostasis, and platelet activation, emphasizing their relevance to the study.

The presence of free intraperitoneal fluid collected inside the Pouch of Douglas was consistent with previous literature ^46^. In individuals with endometriosis, free fluid was detected in 16% of the patients. A considerable number of samples (45%) were contaminated with blood and were thus excluded from this investigation. The single-particle phenotyping investigation on small EVs indicated the presence of platelet-derived GPIIb/IIIa and PF4 in all tested samples, which is consistent with the bioinformatic analysis highlighting platelet degranulation amongst the enriched pathways in this microenvironment. On the other hand, the assessment of the ratio of this particular EV subset revealed significantly increased values in individuals who had been diagnosed with endometriosis.

The release of follicular fluid during ovulation seems to influence the composition of the peritoneal fluid ^47^. Moreover, clinical and experimental data suggest platelets regulate hypothalamic–pituitary–ovarian axis functions ^48^. To evaluate the impact of the ovulatory phase on our findings, we quantified the proportion of platelet-derived small EVs in this biological fluid. The isolation process from follicular fluids leads to the production of a population of small EVs. The percentage of platelet-derived EVs in the investigated samples was less than 3%, a substantially lower amount than that observed in the peritoneal fluids of endometriosis patients. Nevertheless, this observed proportion was similar to the levels identified in the control samples. Thus, despite acknowledging the potential influence of follicular fluid on peritoneal fluid composition, the results of this initial investigation suggest no significant impact on the extent of the platelet-derived subpopulation of small EVs.

## 5. Conclusion

Platelet activation assessment in the microenvironment of endometriosis represents an essential tool for clinical investigations that aim to understand the role of platelets in this pathology and how their inhibition can have a therapeutic effect. The pilot study findings indicated an increase in platelet activity in the peritoneal fluid of patients diagnosed with endometriosis, and emphasize the potential of single-particle phenotyping analysis of the small EVs based on platelet biomarkers. Additional studies are essential to elucidate the precise contribution of platelet activity to endometriosis’s pathogenesis and assess its viability as a biomarker for disease diagnosis and monitoring.

## Author contributions

BB: Conceptualization, Investigation, Methodology, Formal Analysis. RDF: Methodology, Formal Analysis. SM: Methodology, Formal Analysis. BP: Methodology, Formal analysis. RL: Methodology, Formal analysis. FV: Investigation, Methodology, Formal Analysis. MB: Investigation, Methodology, Formal Analysis. AM: Methodology, Formal Analysis. GDL, FR, GZ, FZ, GR: Provision of study materials and patients. VC: Methodology, Formal Analysis. BB: Conceptualization, Investigation, Methodology, Formal Analysis. SB: Conceptualization, Investigation, Methodology, Formal Analysis, Supervision, Writing – original draft and funding acquisition. All authors have read and agreed to the published version of the manuscript.

## Ethics approval and consent to participate

The experimental protocol was approved by the Institutional Review Board (IRB-BURLO 01/2022, 09.02.2022) and patients were asked to sign an informed consent form.

## Data Availability

The data supporting the findings of this study are available in the open dissemination research data repository Zenodo with the digital object identifier (DOI): 10.5281/zenodo.10849687.

