## Supplementary 1 for "Platelets as key cells in endometriosis patients: insights from small extracellular vesicles in peritoneal fluid and endometriotic lesions analysis"

### Control secretive phase

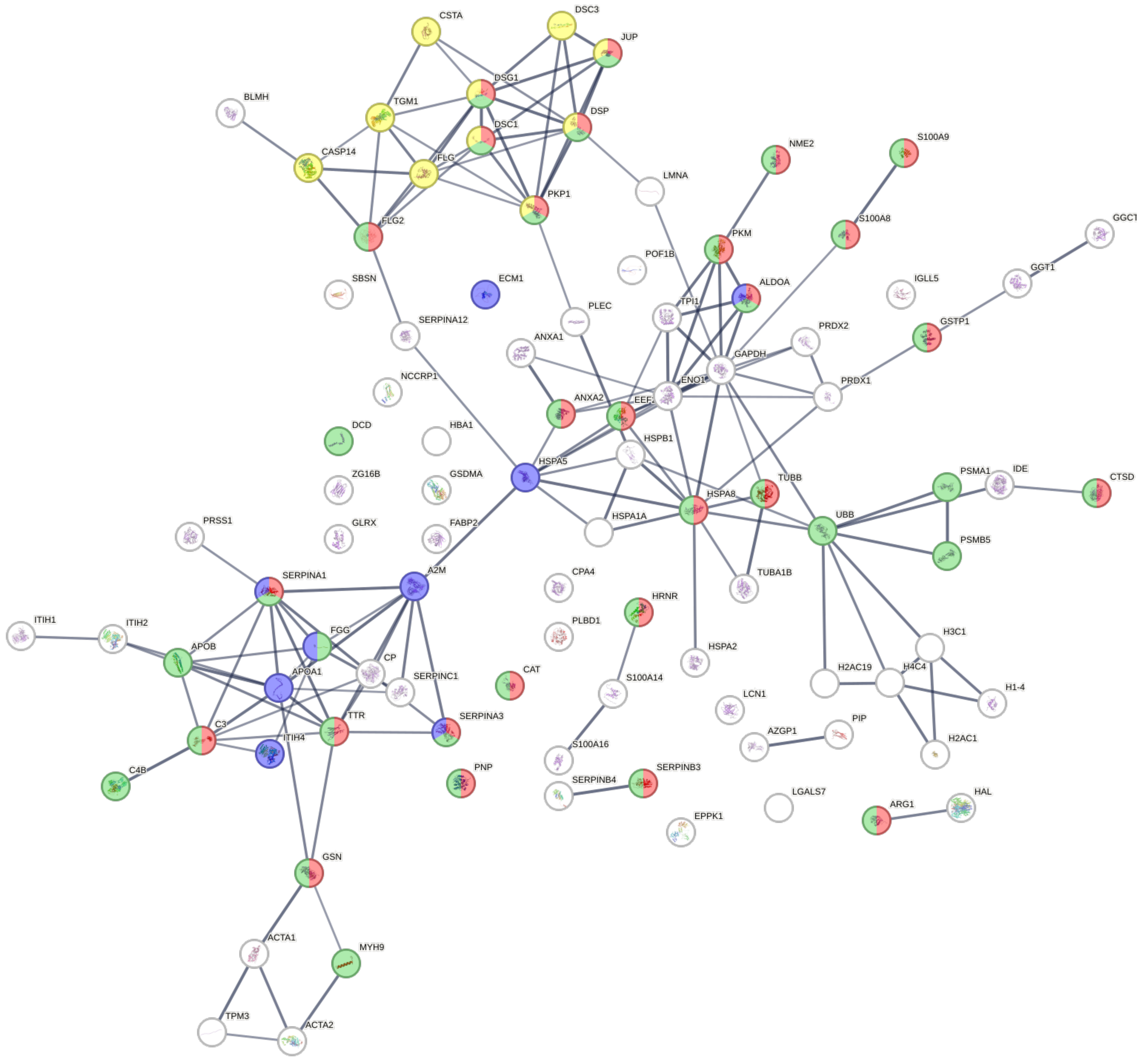


| *Reactome pathways* | *FDR* | *Observed gene* | *Color code* |
|---|---|---|---|
| Neutrophil degranulation | 7.62e-18 | 27 | 🔴 |
| Innate Immune System | 1.87e-17 | 35 | 🟢 |
| Immune System | 2.63e-12 | 39 | |
| Formation of the cornified envelope | 6.80e-07 | 10 | 🟡 |
| Apoptotic execution phase | 5.96e-06 | 7 | |
| Apoptosis | 6.65e-06 | 10 | |
| Platelet degranulation | 6.65e-06 | 9 | 🔵 |
| Apoptotic cleavage of cellular proteins | 1.61e-05 | 6 | |
| Post-translational protein phosphorylation | 1.83e-05 | 8 | |
| Regulation of Insulin-like Growth Factor (IGF) | 4.14e-05 | 8 | |

### Endometriosis stage I/II secretive phase

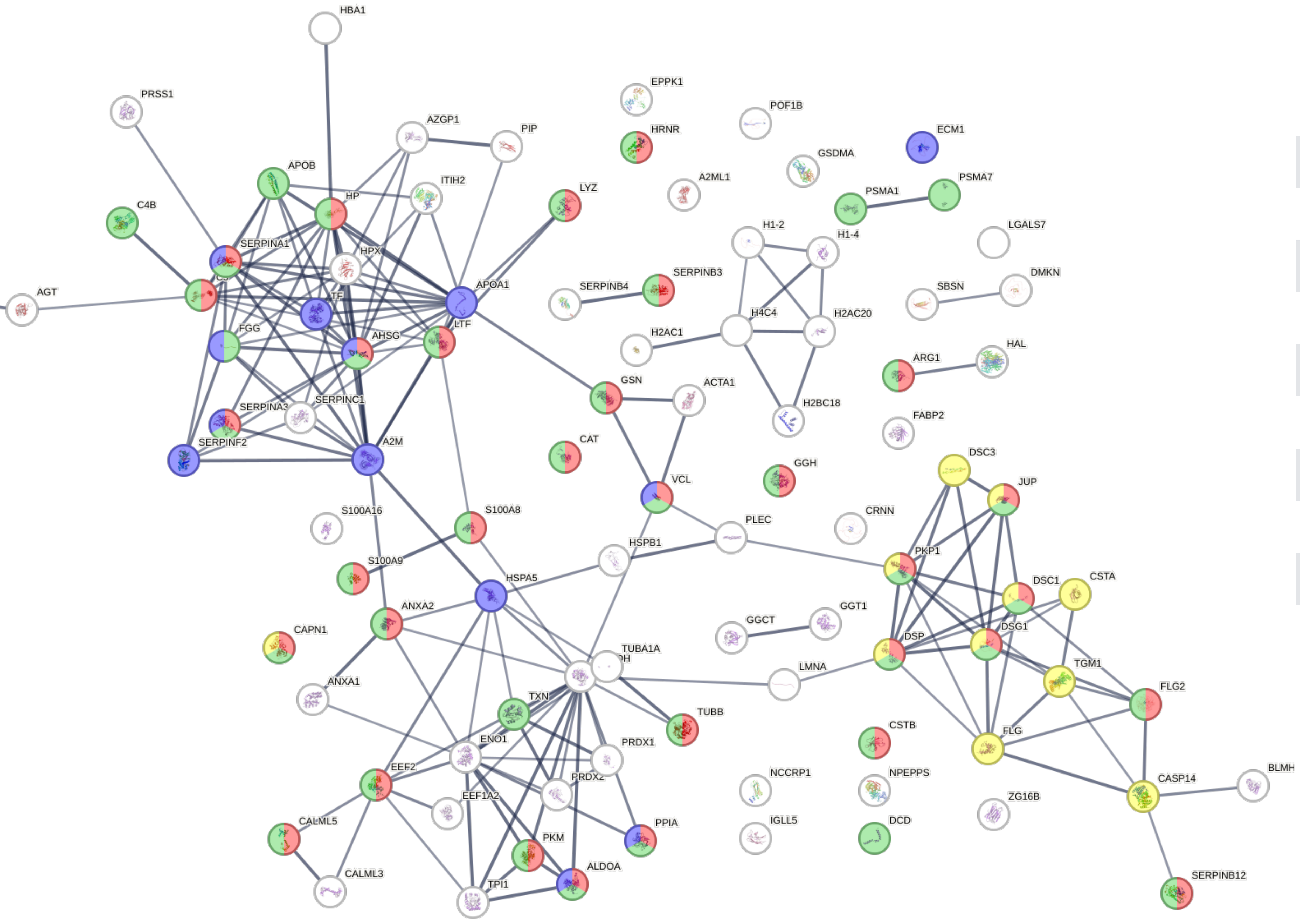


| *Reactome pathways* | *FDR* | *Observed gene* | *Color code* |
|---|---|---|---|
| Neutrophil degranulation | 5.34e-25 | 33 | 🔴 |
| Innate Immune System | 2.54e-22 | 40 | 🟢 |
| Immune System | 6.19e-17 | 45 | |
| Platelet degranulation | 9.17e-11 | 13 | 🔵 |
| Formation of the cornified envelope | 3.10e-08 | 11 | 🟡 |
| Apoptotic execution phase | 1.68e-07 | 8 | |
| Post-translational protein phosphorylation | 1.40e-06 | 9 | |
| Hemostasis | 1.93e-06 | 17 | |
| Regulation of Insulin-like Growth Factor (IGF) | 3.50e-06 | 9 | |
| Apoptosis | 4.35e-06 | 10 | |
